# Physical modeling of a sliding clamp mechanism for the spreading of ParB at short genomic distance from bacterial centromere sites

**DOI:** 10.1101/2020.07.22.213413

**Authors:** Jean-Charles Walter, Jérôme Rech, Nils-Ole Walliser, Jérôme Dorignac, Frédéric Geniet, John Palmeri, Andrea Parmeggiani, Jean-Yves Bouet

## Abstract

Bacterial ParB partitioning proteins involved in chromosomes and low-copy-number plasmid segregation have recently been shown to belong to a new class of CTP-dependent molecular switches. Strikingly, CTP binding and hydrolysis was shown to induce a conformational change enabling ParB dimers to switch between an open and a closed conformation. This latter conformation clamps ParB dimers on DNA molecules, allowing their diffusion in one dimension along the DNA. It has been proposed that this novel sliding property may explain the spreading capability of ParB over more than 10-Kb from *parS* centromere sites where ParB is specifically loaded. Here, we modeled such a mechanism as a typical reaction-diffusion system and compared this ‘Clamping & sliding’ model to the ParB DNA binding pattern from high-resolution ChIP-sequencing data. We found that this mechanism cannot account for all the *in vivo* characteristics, especially the long range of ParB binding to DNA. In particular, it predicts a strong effect from the presence of a roadblock on the ParB binding pattern that is not observed in ChIP-seq. Moreover, the rapid assembly kinetics observed *in vivo* after the duplication of *parS* sites is not easily explained by this mechanism. We propose that ‘Clamping & sliding’ might explain the ParB spreading pattern at short distances from *parS* but that another mechanism must apply for ParB recruitment at larger genomic distances.

## Introduction

Faithful segregation of the full set of genetic information is essential for all living cells. In bacteria, segregation follows DNA replication and involves the separation and transportation of the new copies in opposite directions along the longitudinal cell axis (Bouet et al., 2014). Partition of chromosomes and low copy number plasmids mainly relies on ParAB*S* systems (for reviews; Bouet and Funnell, 2019; Jalal and Le, 2020). They encode a Walker-type ATPase (ParA) and a site-specific DNA binding protein (ParB), which binds to *parS* centromere sites. ParA action separates the ParB assemblies nucleated on *parS* sites, which are always located near the origin of replication, and actively relocates them toward opposite cell poles. This process ensures that every daughter cell receives at least one copy of the replicated DNA molecules.

Recent studies on the bacterial partition protein ParBs have revealed a new activity, namely CTP binding (Jalal et al., 2020; Osorio-Valeriano et al., 2019; Soh et al., 2019). ParB proteins thus now emerge as a new class of CTP-dependent molecular switches. ParB dimers bind specifically to a few short *parS* DNA sites, usually 16-bp, which subsequently nucleate the assembly of hundreds of ParB in their spatial vicinity. This leads to the formation of highly concentrated but dynamic nucleoprotein complexes of small size (~40 nm; Guilhas et al., 2020), with ParB binding to non-specific DNA over large genomic distances (> 10-Kb; e.g. Breier and Grossman, 2007; Debaugny et al., 2018; Lagage et al., 2016; Rodionov et al., 1999; Sanchez et al., 2015), and visible as bright foci in fluorescent microscopy (Breier and Grossman, 2007; Diaz et al., 2015, 4694; Erdmann et al., 1999; Lagage et al., 2016; Lim et al., 2014). Biochemical studies in the presence of CTP and the resolution of a ParB-CDP co-crystal structure suggested that a sliding clamp mechanism controlled by CTP binding could be the basis for this clustering of numerous ParBs around the *parS* sequence (Jalal et al., 2020; Soh et al., 2019).

The ParB-DNA nucleoprotein complex is involved in the intracellular positioning of the *parS* centromere sequences and their segregation toward opposite poles of the cell through its interaction with the cognate ParA ATPase (for reviews; Bouet and Funnell, 2019; Jalal and Le, 2020). CTP binding and hydrolysis control ParB activities (Jalal et al., 2020; Osorio-Valeriano et al., 2019; Soh et al., 2019). Apo-ParB dimers adopt an open conformation and bind to DNA (Fig. 1A). Specific binding to *parS* activates CTP binding which induces a conformational change that converts ParB dimers into a closed conformation (gate closure) entrapping double strand DNA. The clamped-ParB favor *parS* unbinding as suggested by (i) the steric clash for *parS* binding modeled in the CDP-bound ParB crystal structure (Soh et al., 2019) and (ii) the ParB release from *parS* in the presence of CTP (Jalal et al., 2020). Upon release from *parS* sites, clamped-ParB dimers would thus remain trapped along the DNA on either side of *parS* until the gate opens to release ParB from the DNA (Fig. 1B). Soh et al. (2019) thus proposed that a ParB sliding clamp mechanism, restricting ParB around *parS* sites, might give rise to the characteristic ParB DNA binding distribution observed by ChIP assays (e.g. Breier and Grossman, 2007; Rodionov et al., 1999; Sanchez et al., 2015). Importantly, CTP binding and hydrolysis by ParB proteins was demonstrated for plasmid- and chromosomally-encoded ParABS systems (Soh et al., 2019).

**Figure 1:**
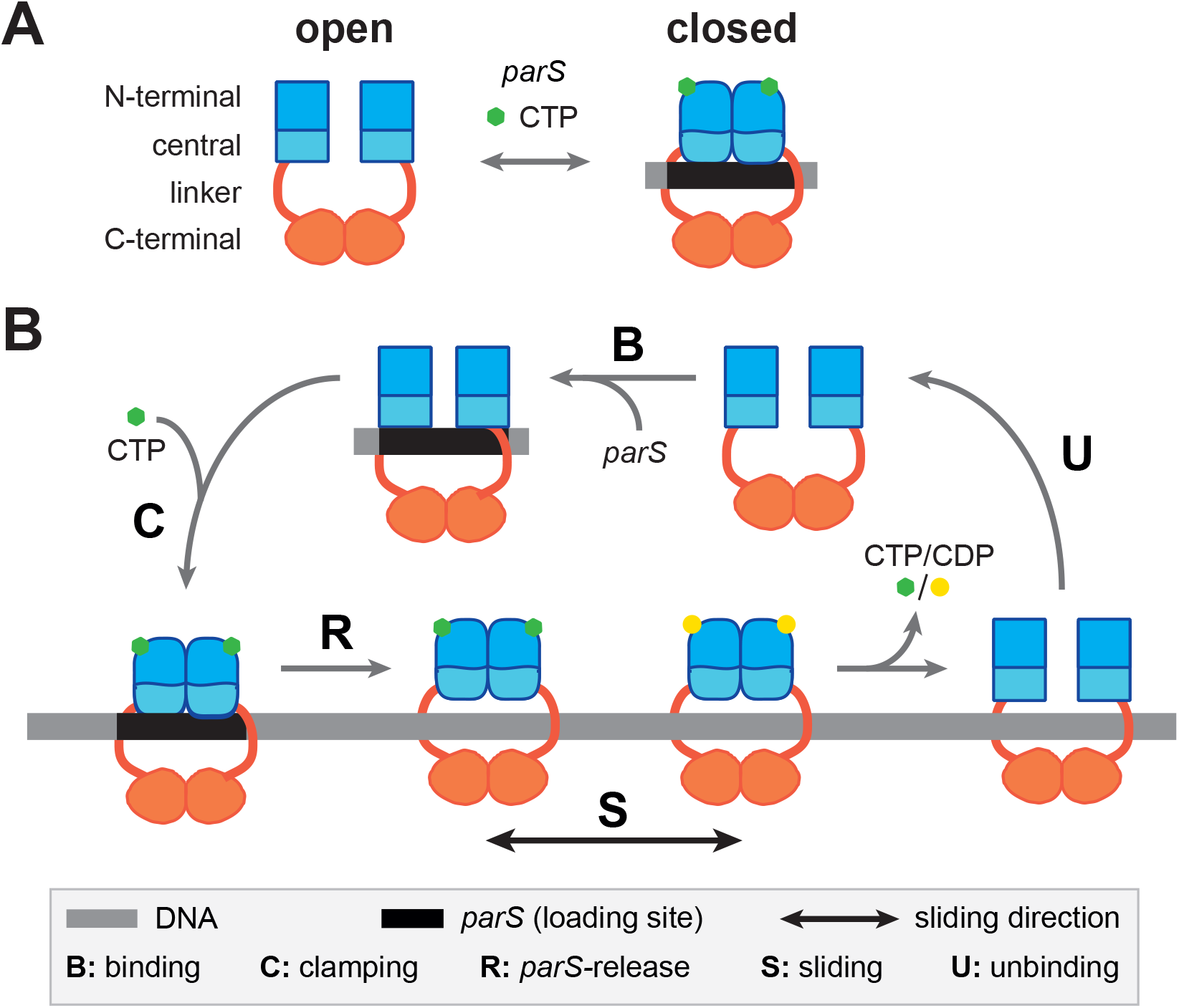
Schematic cycle for ParB clamping and sliding by diffusion along DNA. **A-** Open and closed conformations mediated by CTP and *parS* DNA. ParB is a homodimer composed of a C-terminal dimerization domain (orange) link to the central (light blue) and N-terminal (dark blue) domains by a flexible linker (red). The central domain contains the two DNA binding motifs for *parS* binding (Sanchez et al., 2013). The N-terminal part contains the ParA interaction domain, the arginine-like motif, the CTP binding motif and the multimerization domain (Ah-Seng et al., 2009; Soh et al., 2019; Surtees and Funnell, 1999). In the presence of *parS* and CTP, ParB dimer forms a clamp around the DNA. **B-**Schematic representation for the ‘Clamping & sliding’ model displaying the five key steps. The open conformation of ParB dimer enables DNA binding. Upon specific binding to *parS* centromere (step B), ParB undergoes a conformational change promoting CTP binding which subsequently induces ParB to form a clamp around *parS* (step C). Clamping promotes its release from *parS* allowing the ParB clamp to slide away from *parS* by diffusion (step S), and to free the *parS* loading site for next round of loading. The parameters used in the physical modeling *B*, *R*, *D* and *U* correspond to the *parS* binding (B), ParB clamping (C) and release (R), free diffusion during sliding (S), and DNA unbinding (U) steps, respectively. Note that (i) *R* is the total release rate from *parS* which is, to a first approximation, equal on both sides of *parS* (not represented on the schematic representation for simplicity), therefore ParB clamps are loaded on each side at a rate *R*/2, and (ii) the stage at which CTP hydrolysis occurs was not determined in the original clamping models (Jalal et al., 2020; Soh et al., 2019).

ParB was found to cluster around *parS* based (i) on gene silencing in the vicinity of *parS* (Lynch and Wang, 1995; Rodionov et al., 1999), (ii) on ParB binding over more than 10-Kb on both sides of *parS* (Murray et al., 2006; Rodionov et al., 1999; Sanchez et al., 2015), and (iii) on the lower accessibility of proteins around *parS* DNA (Bouet and Lane, 2009; Lynch and Wang, 1995). However, the quantity of ParB dimers present in the cell is insufficient to continuously cover the observed spreading zone. Several groups have carried out physical modeling studies (Broedersz et al., 2014; Debaugny et al., 2018; Sanchez et al., 2015; Walter et al., 2020; Walter et al., 2018) to investigate the mechanisms that could account for the assembly of ParB (reviewed in Funnell, 2016; Jalal and Le, 2020). By testing the ‘1D-spreading’, ‘Spreading & bridging’ in the strong coupling limit, ‘Looping & clustering’ and ‘Nucleation & caging’ models on high-resolution ChIP-sequencing data, only the latter model based on low affinity but synergistic interactions actually fits best observations (Debaugny et al., 2018). However, all these models were proposed before the finding that ParB can clamp and diffuse unidimensionally along the DNA.

Here, we developed a physical modeling approach based on the newly proposed sliding mechanism, namely ‘Clamping & sliding’, involving the loading of ParB at *parS* sites with subsequent free diffusion as a protein clamp along the DNA track, followed by its unbinding upon clamp opening (Fig. 1B). We then compared this novel physical model with the ParB DNA binding pattern from high-resolution ChIP-sequencing data reported for the ParAB*S* system of the F plasmid. Using recent estimations of the release and unbinding parameters (Jalal et al., 2020) and of the diffusion coefficient (Guilhas et al., 2020), we showed on the basis of a physical analysis with biological parameters of incoming and outgoing flux of ParB on DNA that ‘Clamping & sliding’ is not able to account for the number of particles found *in vivo* on the F plasmid, especially at large distances from *parS*. Importantly, one-dimensional translocation is expected to be interrupted by obstacles on DNA. The modeling of the effect of a roadblock failed to describe the ChIP-seq pattern obtained in the presence of a natural roadblock. Moreover, imaging ParB clusters using epifluorescence microscopy at high temporal resolution indicated a rapid recovery of ParB fluorescence intensity at the onset of DNA replication in contrast to the model prediction. These results suggest that the ‘Clamping & sliding’ model can only partly account for the assembly mechanism. We thus propose that a ‘Clamping & sliding’ mechanism might explain the ParB spreading pattern only at short distances from *parS* but that another mechanism must apply for ParB binding at long distance.

### The ‘Clamping & sliding’ model

The DNA is modeled as a circular filament of length *L* displaying *N* discrete sites of 16-bp (footprint of a ParB dimer; Bouet and Lane, 2009). ParB dimers (i) bind specifically to *parS*, (ii) switch conformation and bind CTP, (iii) convert to protein clamps, (iv) release *parS*, (v) freely diffuse on DNA, and (vi) unbind DNA by opening the clamp (Fig. 1B). Therefore, ParB can only enter the filament by a unique loading gate, *parS*, at the binding rate *B* (Fig. 1B). It is released from *parS* at the rate *R*, and it can diffuse unidimensionally on the DNA, with a diffusion coefficient *D*, but cannot cross *parS*, which is always assumed to be occupied by ParB. Lastly, ParB clamps exit anywhere from the circular DNA filament by opening the clamp at the unbinding rate *U*. We thus modeled the ‘Clamping & sliding’ mechanism by a typical reaction-diffusion model. Notably, in contrast to all other models at thermodynamic equilibrium previously proposed (see above), this steady-state sliding mechanism is out-of-equilibrium. Indeed, the binding and subsequent release of ParB at *parS* only creates an oriented gradient of proteins leading to a stationary, oriented flux of ParBs. Thus, the system is brought out-of-equilibrium due to these particular boundary conditions. Note that in the framework of the ‘Clamping & sliding’ model, ParB non-specific DNA binding activity is not relevant due to the high concentration of competitor DNA represented by the nucleoid.

In the cells, ParB dimers can be found in two states: (i) assembled in clusters around *parS*, which represents over 90% of ParB present in the cell and (ii) freely diffusing over the nucleoid corresponding to the remaining ParB population, which acts as a reservoir for the clusters (Guilhas et al., 2020; Sanchez et al., 2015). These two populations may represent the sliding clamps and the free ParBs, respectively. Therefore, in the stationary regime, the flux balance of ParB in the partition complex is *J*_*in*_ = *J*_*out*_, with the current of incoming particles *J*_*in*_ = *R*, and outgoing particles *J*_*out*_ = *N*_*a*_*U*, where *N*_*a*_ is the number of particles clamped to DNA. This leads to the conservation of the average number *N*_*a*_ of clamped-ParB, imposed by the two rates, *R* and *U*:

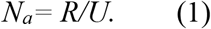

The time-dependent density *ρ*(*x*, *t*) of ParB can be described by the following equation:

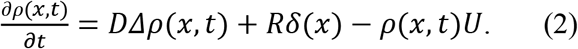

The stationary solution for a coarse grained number density, *n*(x)=*ρ*(*x*)*δx*, is readily obtained from the steady-state solution of Eq.(2) as:

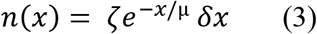

where *δx* = 16-bp corresponds to one footprint of ParB, 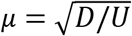 is the characteristic length scale (in bp) of the decay corresponding to the typical distance performed by ParB by diffusion before detachment, and 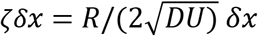 is the overall amplitude of the density decay, which linearly increases with the release rate *R*. In the next section, we applied this formalism using biologically determined parameter values.

## Results

### Simulation of the “Clamping & sliding’ model using biological parameters for ParB

Our physical modeling for the ‘Clamping & sliding’ mechanism contains four kinetic parameters (see Fig. 1B). We estimated these parameters as follows: (i) We considered that *parS* is always occupied by a ParB dimer. Indeed, the ParB binding to *parS* depends on the population of ParB that is free to diffuse over the nucleoid. The freely diffusing ParB concentration was estimated at ~0.14 μM and ~0.48 μM for ParB from plasmid F (ParB_F_) and from the chromosome of *C. crescentus* (ParB_*Ccre*_) (see Table S1; Bouet et al., 2005; Lim et al., 2014), well above the *K*_*D*_ for ParB_F_-*parS*_F_ interaction (~2 nM; Ah-Seng et al., 2009) and chromosomal ParB-*parS* (~20-50 nM; Fisher et al., 2017). In addition, the ParB_F_ binding rate, *B*_F_, with an apparent association constant *K*_*on*_ ~2 ×10^5^ mol^−1^ s^−1^ (Ah-Seng et al., 2009) is several orders of magnitude higher than the other rates used in the model (see below). The ParB-*parS* binding step is therefore not limiting; (ii) the apparent diffusion coefficient of ParB_F_ clustered around *parS*_F_, *D*_F_ ~ 0.05 μm^2^ s^−1^ (or ~ 4.3×10^5^ bp^2^ s^−1^) as measured by super-resolution microscopy (Guilhas et al., 2020); (iii) the release of ParB from *parS*, *R*_*Ccre*_ ~ 0.1 s^−1^ (Fig. S1A-B); and (iv) the unbinding of the ParB clamp from non-specific DNA *U*_*Ccre*_ ~ 0.017 s^−1^ (Fig. S1A,C). These two latter parameters were obtained from the *in vitro* biochemical experiments, performed with ParB_*Ccre*_ in the presence of CTP (Fig. 2A in Jalal et al., 2020), by calculating the kinetic rates using nonlinear regression fits (see Fig. S1 and Methods). Eq.(2) contains only the three kinetic parameters *D*, *R* and *U*. We accounted for the assumption that the *parS* site is always occupied by a ParB with a constant source term, i.e. a delta-function *δ*(*x*).

**Figure 2:**
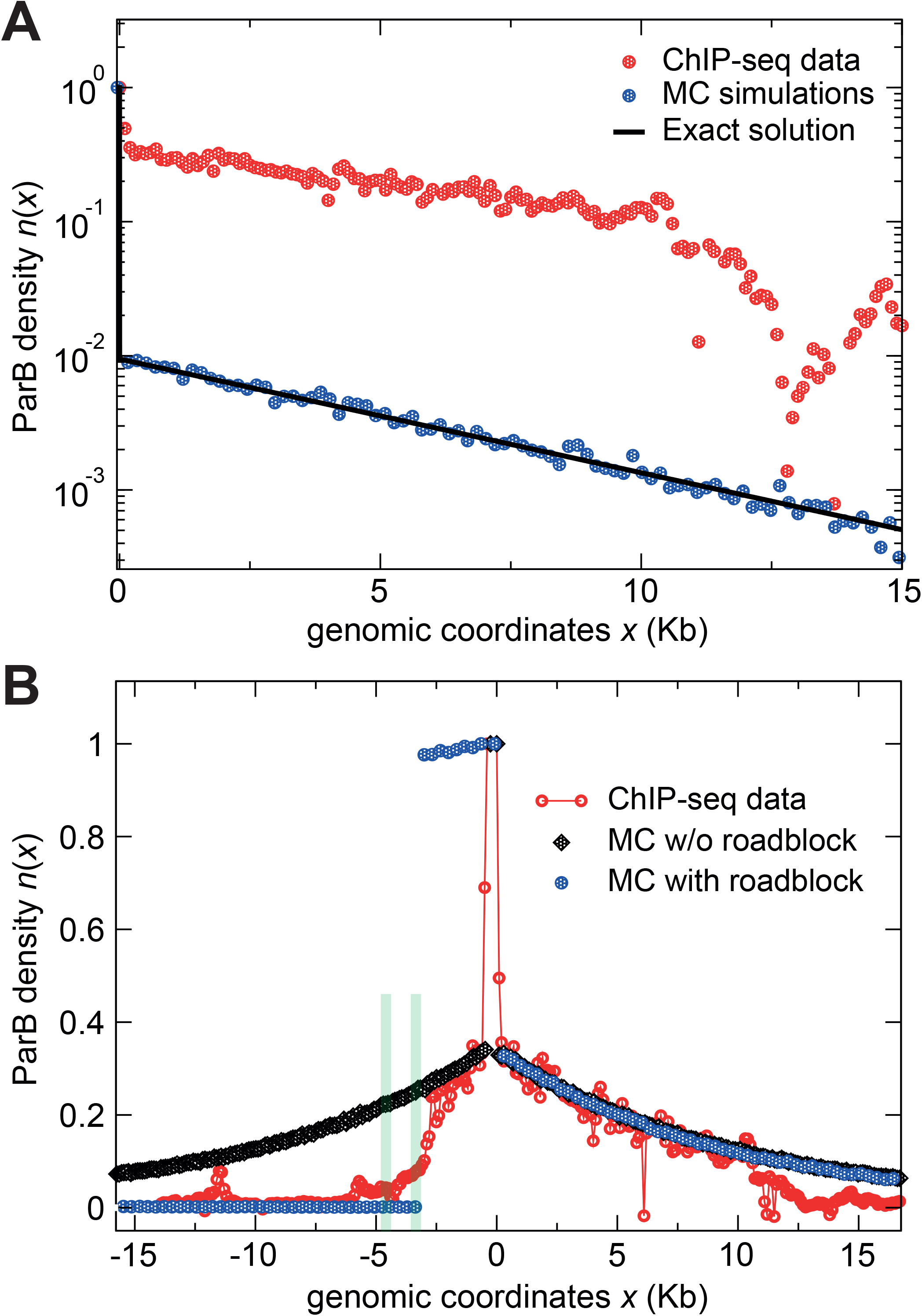
Comparisons between the ‘Clamping & sliding’ model and the ParB_F_ DNA binding pattern. High-resolution ChIP-sequencing data from a previous study (Sanchez et al., 2015) display the average number of reads per 100-bp windows as a function of the genomic coordinates (red circles). Note that the signal at *parS*_F_ (coordinate 0) is normalized to 1 by averaging the number of reads over the 550-bp centromere locus. **A-** The model from Eq.(2) contains the three kinetic parameters *D* = 4.3×10^5^ bp^2^ s^−1^ (Guilhas et al., 2020), *R* = 0.1 s^−1^ and *U* = 0.017 s^−1^ (Fig. S1B-D). The exact solution (black line) and the Monte Carlo (MC; blue dots) simulation data are plotted on the right side of *parS*_F_. The theoretical prediction is compared to the ChIP-seq data (Sanchez et al., 2015). The large discrepancy between the model and the ChIP-seq data is essentially due to the amplitude (value of *R*). **B-** Simulation in the absence (black diamonds) and in the presence (blue circles) of a roadblock compared to the Chip-seq data with the optimized parameters (*R* = 1.91 s^−1^ and *U* = 0.0047 s^−1^). The two strong roadblocks (green bars), present at ~3-kb and 4.5-kb on the left of *parS*_F_, are due to the binding of RepE to *incC* and *ori2* iterons, respectively (Rodionov et al., 1999; Sanchez et al., 2015). Note that the two MC simulations overlap on the right side of *parS*_F_.

We introduced these parameter values in our physical model to estimate the number of attached particles, and we obtained, from Eq.(1), an average number of clamped ParB dimers per DNA molecule *N*_*a*_ = *R/U* ~6. This value is much lower than the *N*_*a*_ ~250 or ~200 (see Table S1) observed experimentally for ParB_F_ and ParB_*Ccre*_, respectively (Adachi et al., 2006; Bouet et al., 2005; Lim et al., 2014) indicating that the biological values for *R* and/or *U* cannot for the number of ParB per cluster. Despite this discrepancy, we further plotted, in Fig. 2A, the theoretical prediction for the profile given by Eq. (3) and, as a benchmark, the corresponding Monte Carlo simulations. These simulations consist of a discrete version of the ‘Clamping & sliding’ model applied on a DNA molecule of length *L* = 60-Kb, which corresponds to the actual size of the F plasmid used for the ChIP-sequencing data (see Methods), giving *N* = 3750 non-overlapping ParB sites. From Eq. (3), the exponential decay has a characteristic length μ ~5.1-Kb and an amplitude *ζ* ~6×10^−4^*bp*^−1^, leading to the coarse-grained amplitude *ζδx* ~10^−2^. As shown in Fig. 2A, analytic and numerical predictions are both in excellent agreement with each other but result in a very low density profile around *parS*, unable by far to describe the ChIP-seq profile (red symbols). *N*_*a*_, the total number of ParB on DNA, corresponds to the integral of *n(x)*. Thus, a low value of *N*_*a*_ ~6 implies directly a low value of the overall amplitude of *n(x)* with the biological parameters from Fig. S1 compared to the ChIP-seq data. Also, the slopes of the decay are different: the ChIP-seq profile displays a slower decay, suggesting a larger characteristic length μ (discussed in the next paragraph). Since the parameters arise from *in vitro* assays performed on a small and constrained DNA molecule (Jalal et al., 2020), the value of *U*_*Ccre*_ may be overestimated compared to *in vivo* conditions (see discussion).

Another approach is to fit directly the *U* and *R* parameters, the DNA unbinding and *parS* release rates, respectively, from the *in vivo* ParB DNA binding profile. We estimated these two values (*D* remains the same) leading to the best fit to the ChIP-seq profile. In Fig. 2B, the ChIP-seq data (red symbols) display an enrichment at *parS*_F_ since these sites are saturated due to the high value of the binding rate *B*_F_. The drop on both ends of *parS*_F_ and the subsequent slower decay can be interpreted in the framework of the ‘Clamping & sliding’ model described by Eq.(3). We fitted the right side of the ChIP-seq profile (in the range 0 < *x* < 11-Kb) because the decay is longer with no roadblock hindering ParB binding, compared to the left side. We used the functional form *n*(*x*)= *Ae*^*-Bx*^ and first fitted the parameter *B* = 1.04 (+/− 0.1)×10^−4^ bp^−1^, which is related to μ in Eq.(3) as 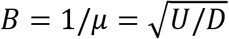. Assuming D = 4.3×10^5^ bp^2^ s^−1^, we obtained *U* = 4.7 (+/− 0.5) ×10^−3^s^−1^. This fitted value is ~4-times lower than the *in vitro* estimate from Fig. S1C. Also, note that the optimized value of μ ~9.6-Kb is ~2-times higher than the value estimated in the previous paragraph with the *in vitro* parameters (Fig.2A). Second, we fitted *A* = 0.34 (+/− 0.1), which is related to ζ in Eq.(3) as 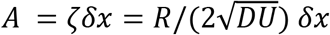. Assuming the previous values of *D* and *U*, this leads to *R* = 1.91 (+/− 0.05) s^−1^, a value ~20-times higher than the *in vitro* one. This theoretical prediction provides a symmetrical ParB DNA binding pattern with respect to *parS* (black diamonds, Fig. 2B), which also fits with the decrease observed experimentally on the left side of *parS*_F_ up to ~3-Kb, beyond which the RepE/*IncC* protein-DNA complexes interfere with ParB_F_ binding (see below).

Using these optimized values, the number of particles *N*_*a*_ clamped to DNA at the steady-state is *N*_*a*_ = *R/U* ~400, which is roughly compatible with the estimation of ~250 or ~200 in average per cluster at *parS* for ParB_F_ or ParB_*Ccre*_, respectively (Table S1). Also, these values are, to a first approximation, within the biological range of ParB release time from *parS* τ_*R*_ ~1/*R* ~0.5 sec and ParB unbinding time from the DNA τ_*U*_ ~1/*U* ~213 sec.

### Modeling the ParB binding pattern in the presence of roadblock

The ParB binding pattern is strongly impaired by protein-DNA complexes acting as roadblocks which strongly reduce the ParB density on DNA beyond these sites as observed both *in vivo* and *in vitro* (Breier and Grossman, 2007; Jalal et al., 2020; Rodionov et al., 1999; Sanchez et al., 2015). In order to test the prediction of roadblocks in the ‘Clamping and sliding’ model, we simulated the effect of a protein-DNA complex present at ~3-Kb on the left side of *parS* which mimics the position of the RepE-*IncC* roadblock present on the plasmid F (Sanchez et al., 2015). This roadblock is modeled by a no-flux boundary condition, *i.e.* particles cannot cross the barrier. By using the fitted parameters adapted for the case without roadblock (see previous section), we obtained the curve presented in Fig. 2B (blue circles). The symmetry between the left and right sides is clearly broken with ParB particles accumulating between *parS* and the roadblock as shown by the formation of a plateau starting immediately after the *parS* site. The ParB that are released on the left side of *parS* diffuse on an isolated domain of DNA of ~3-Kb comprised between the two diffusion barriers constituted by *parS* and the roadblock. ParB dimers only detach from the DNA segment at the unbinding rate *U.* Given that ParB diffuses with a diffusion coefficient *D* = 4.3×10^5^ bp^2^ s^-1^, the size of the isolated domain is covered in a few seconds. In a first approximation, if we consider that ParB is uniformly released from DNA between *parS* and the roadblock, we conclude that *R*/(2*U*) ~200 ParB are homogeneously distributed over 3-Kb, giving a nearly saturated average density ~1. The modeling is thus in clear contrast with the observed data from ChIP-sequencing showing that (i) the ParB binding pattern is nearly symmetrical over 3-Kb on both sides of *parS*_F_ and (ii) no plateau is observed between *parS* and the roadblock. This finding may also argue for an unbinding rate higher than the model prediction.

We reasoned that the unbinding rate could be higher when encountering a roadblock. Indeed, we found that the ParB unbinding kinetics is best fit with a two phase exponential decay giving two unbinding rates (Fig. S1C-D). We used the lowest rate (*K*_*off_slow*_ = 0.017 s^−1^) in our initial modeling since it may best correspond to the physiological condition on large *parS* DNA molecules. However, ParBs may reach saturation in front of a roadblock, as it occurs *in vitro* on the small DNA probe with fixed ends (see legend Fig. S1C), leading to an initial fast unbinding rate (*K*_*off_fast*_ = 0.086 s^-1^). We therefore tested the higher rate *U*_*Ccre*_ = 0.086 s^−1^ on the left side of *parS* in front of the roadblock. With this value, 20-times higher than on the right side of *parS*_F_, the pattern is not saturated but still displayed a plateau (Fig. S2). This contrasts with the ChIP-seq data, which clearly indicates that the ‘Clamping & Sliding’ mechanism is not able to account for a barrier to ParB clamp diffusion. Moreover, this data points out that sliding by free, unidimensional diffusion as a clamp would be highly dependent on any obstacle present along the DNA. These obstacles, mainly proteins stably bound to DNA, would strongly impair ParB sliding by reducing its diffusion away from *parS*. Notably, ChIP-seq data present ParB DNA binding patterns with dips and peaks, *i.e.* indicating numerous obstacles along the DNA (Fig. 2B, and e.g. Baek et al., 2014; Bohm et al., 2020; Breier and Grossman, 2007; Debaugny et al., 2018; Lagage et al., 2016). Such a pattern indicating a recovery of the ParB binding signal after strong dips could not be easily explained by the sliding clamp mechanism (see discussion).

Based on these observations, we thus conclude that the ‘Clamping & sliding’ model does not apply over a large range of genomic coordinates, and thus could not account fully for the assembly of the partition complex. In the following sections, we will only consider the value of the parameters *R* and *U* obtained *in vitro* from ParB_*Ccre*_.

### ParB clusters reassemble faster than predicted by ‘Clamping & sliding’

A ‘Clamping & sliding’ based-mechanism is expected to have an important effect on the *de novo* assembly kinetics of ParB. In this model, the ParB dimers load sequentially from a unique source, *parS* (Fig. 1B), thus their accumulation in clusters should increase progressively at the rate *R* up to a stationary population average. ParB are considered in steady state within the clusters, as suggested by fluorescence microscopy and fluorescence recovery after photobleaching (FRAP) experiments (Debaugny et al., 2018; Guilhas et al., 2020), except when the centromere site is replicated. DNA replication is highly processive, with a rate of DNA unwinding and duplication of ~1-Kb per second (Kelman and O’Donnell, 1995). ParB clamped at and around *parS* should therefore be unloaded or dispersed very rapidly when DNA polymerase III holoenzyme crosses over the ParB binding zone. ParB binding to newly duplicated *parS* sites is fast but its release as a sliding clamp on either side is limited by the ParB release rate *R* ~ 0.1 s^−1^ (see note in Figure S1B). In the framework of the ‘Clamping & sliding’ model, it would take ~20-40 min to load an average stationary population of ~125-250 ParB after the passage of the replication fork. We argue that, at the onset of replication, such disassembly/reassembly of ParB should be visible in time-lapse epi-fluorescence microscopy.

To test this expectation, we recorded ParB clusters in an *E. coli* strain carrying a 11.1-Kb mini-F plasmid expressing the fully functional ParB_F_-mTq2 fusion protein (Diaz et al., 2015). Images were taken every 5 sec for a period of 10 min and displayed as kymographs (Fig. 3A). Since the generation time in our growth condition is ~100 min, only a few cells would undergo mini-F replication during the recorded period. Note that mini-F replication occurs within 10 sec, and about once per cell cycle and per plasmid (present at ~two copies per chromosome; Frame and Bishop, 1971). In addition, subsequent DNA segregation, visible as ParB_F_ foci splitting, occurs within less than five min after replication (Onogi et al., 2002). In spite of these difficulties, we were able to observe three trajectories (labelled #1, 2 & 5; Fig. 3A) displaying ParB foci more than 5 min before their splitting (indicated by arrows). We observed that (i) the ParB fluorescence intensity is maximal just before the splitting event and lasts at this level between one to three minutes, (ii) the ParB signals do not decrease importantly over the time course recorded. Only a few frames display transiently low signals (indicated by black circles) that last less than two frames (10 sec). However, we could not exclude that these transient faint signals were due to out of focus variation of the ParB foci since such decreases were also observed independently of a splitting event as observed for trajectories #3 and #4 (indicated by white circles).

**Figure 3:**
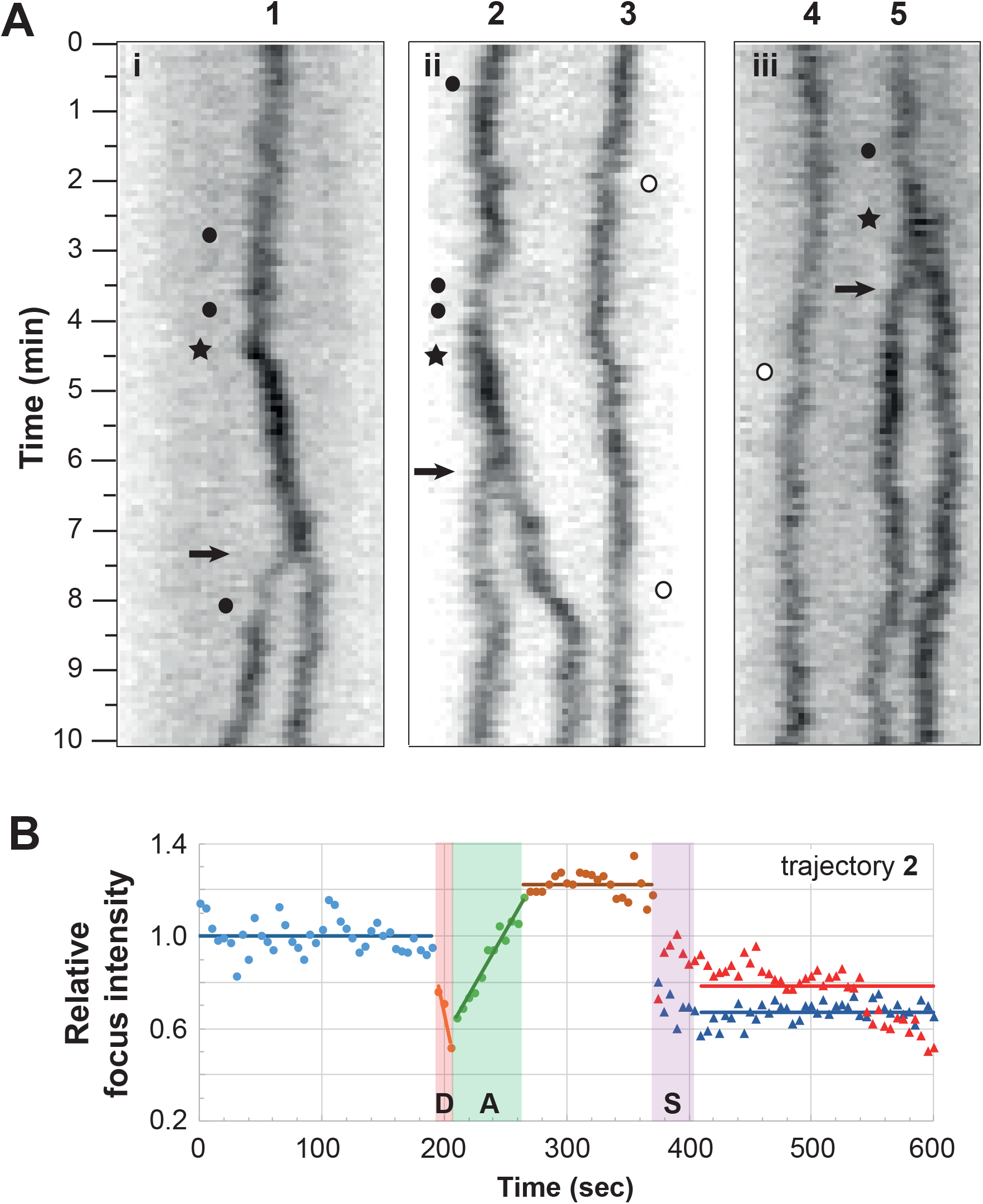
ParB foci do not disassemble for a long time before splitting events. **A-** The trajectories of the ParB_F_-mTq2 foci, labelled 1 to 5, positioned over the nucleoid length, were obtained by kymographs analyses. *E. coli* cells (i-iii), carrying a mini-F plasmid (pJYB249) with its endogenous ParABS_F_ system, expressing a ParB_F_-mTq2 functional fusion protein, were observed by time-lapse epi-fluorescence microscopy. Images were collected every 5 sec over 10 min and converted to kymograph (see Methods). The black arrows indicate the splitting events of ParB foci. Note that the ParB foci intensity increases strongly (black star) 1.5 to 3 minutes before the splitting events. Closed and open circles indicate low ParB fluorescence intensities in traces with or without splitting, respectively. The panel width corresponds to the length of the nucleoid. **B-** Intensity of ParB_F_ fluorescent foci before and after the splitting event. The integrated ParB_F_-mTq2 fluorescence (thin line) from trajectory number 2 in panel A is measured every frame and plotted over time with color data points. The signal was subtracted from the average background level, and normalized to 1 from the average intensity before the drop in intensity (blue dots). Dots and triangles represent the intensity of one and two ParB foci before and after the splitting event (S), respectively. R and L represent the putative centromere replication and the ParB loading steps, respectively. The blue, brown, red and blue horizontal bars represent the mean values of the normalized ParB foci intensity of the one focus before R, before S and the 2 foci after S, respectively. The orange and green bars represent the linear regression of the variation in ParB fluorescent intensities. Note that replication (R) is inferred from the increase in foci intensity that (i) occurs between the 2 plateau of ParB fluorescence mean intensity and (ii) takes place less than 5 min before splitting (S).

To gain quantitative insights, we measured and plotted the relative fluorescence intensity signal within ParB foci over time (Fig. 3B and Fig. S3). We found that it displayed two clear transitions between three stable states: (i) the fluorescence intensity from the ParB focus is maximal before the splitting event (S) and is then nearly equally divided between the two newly formed foci; (ii) less than 5 min before the splitting event, the fluorescence intensity decreased abruptly (D) over 10-15 sec, followed by an increase (A) over a short period of time (20-50 sec). We propose that this decrease might correspond to the passage of the replication fork over *parS* removing ParB from the DNA, and that the immediate increase might correspond to the loading of ParB at *parS* for *de novo* assembly of the partition complexes. Note that, at this stage, a doubling of the ParB fluorescence intensity is not expected since most of ParBs were already present in foci before duplication (Lim et al., 2014; Sanchez et al., 2015). We found that the observed experimental loading time (<60 sec) is much shorter than the time predicted by the ‘Clamping & sliding’ model (~20-40 min; see above). Rather, it is of the same order of magnitude as the time needed for two foci to exchange their ParB (~90 sec; see Debaugny et al., 2018; Guilhas et al., 2020), confirming the fast dynamics of ParB within clusters. This result suggests that the disassembly of the partition complexes is carried out too rapidly to be explained by the ‘Clamping & sliding’ model.

## Discussion

The understanding of the molecular mechanism responsible for the assembly of the bacterial partition complex has resisted three decades of biochemical and molecular studies performed on several ParAB*S* systems from chromosomes and low-copy-number plasmids. How a few ParB molecules bound to *parS* sites cluster hundreds of other ParB in the vicinity of *parS* in a self-assembled high molecular weight structure that serves to position and actively segregate DNA molecules remains puzzling. The recent finding that ParB proteins belong to a new class of CTP-dependent molecular switches has opened new avenues of research to decipher this mechanism. In particular, ParB dimers switch from an opened conformation prone to DNA binding to a close conformation (clamp) upon *parS* and CTP binding that enable the release and sliding away from *parS* as a protein clamp (Jalal et al., 2020; Soh et al., 2019). In this work, we propose a physical model, namely ‘Clamping & sliding’, for this newly proposed CTP-based ParB clamping mechanism.

First, we considered that the *parS* binding site is always occupied by ParB as suggested by the high affinity constants in the ~1 to 50 nM range for all ParBs investigated (e.g. Ah-Seng et al., 2009; Funnell and Gagnier, 1993; Taylor et al., 2015). Therefore, clamped-ParBs must slide away from *parS* by free diffusion since (i) they could not cross back over this strong binding site and (ii) they are pushed away by the successive loading at *parS*.

Second, the ‘Clamping & sliding’ model contains three kinetic parameters (see Eq.(2)), which were inferred from biochemical, microscopy and ChIP-seq analyses. The apparent diffusion coefficient, *D*, was estimated from single molecule live imaging assays (Guilhas et al., 2020). ParB_F_ displays two distinct diffusion modes depending on whether it belongs to a mobile or a clustered fraction. In the mobile fraction (not positioned at *parS* sites), *D* ~ 0.7 μm^2^ s^-1^ which typically corresponds to the diffusion of non-specific DNA binding proteins (Garza de Leon et al., 2017; Stracy et al., 2016; Stracy and Kapanidis, 2017). In the fraction clustered around the *parS*_F_ site, *D* was estimated at ~ 0.05 μm^2^ s^−1^ (Guilhas et al., 2020). We used this latter estimate for *D*_F_ as it corresponds to the ParB_F_ (~ 95%) present in the vicinity of *ParS*_F_ while the former corresponds to the ParB_F_ that are freely diffusing over the nucleoid. Here, we only considered that ParB clamps move unidimensionally and passively by free diffusion along DNA. Indeed, although the role of CTP hydrolysis is not known, it is unlikely that it could provide energy for translocation since (i) the hydrolysis rate is as low as ~40 CTP hydrolyzed per hour, and (ii) ParB still accumulates on DNA in the presence of a non-hydrolysable CTP analog (Jalal et al., 2020; Osorio-Valeriano et al., 2019; Soh et al., 2019).

The release rate of ParB from *parS*, *R*, is the limiting step for creating the flux of ParB clamps on *parS*-proximal DNA. It occurs at the same rate on both sides of *parS* as indicated by all the ChIP assays performed with numerous ParB belonging to plasmids and chromosomes (Baek et al., 2014; Breier and Grossman, 2007; Debaugny et al., 2018; Donczew et al., 2016; Lagage et al., 2016; Rodionov et al., 1999; Sanchez et al., 2015). The value estimated from the *in vitro* data (Fig. S1B and Jalal et al., 2020) is 20-times lower than the best fit from the *in vivo* ChIP-seq data using the ‘Clamping & sliding’ model (Fig. 2B). This difference may arise from the small linear and closed DNA fragment used in the *in vitro* assay (Jalal et al., 2020) compared to the ~60-kb circular DNA for the ChIP-seq. However, the release rate from the former is nicely fitted by a one phase association curve (Fig. S1B) indicating that it is not perturbed by steric hindrance when the DNA probe becomes saturated. This suggests that *R* remains constant from the initial loading stage after replication to the equilibrium state with one ParB clamp released every 20 sec on each side of *parS*. We cannot exclude that the release rate *R* is intrinsic to each ParB from different systems, and that there could be a slight difference between ParB_F_ and ParB_*Ccre*_.

The unloading rate of the clamped-ParB from DNA, *U*_*Ccre*_, was estimated from the *in vitro* unbinding curve (Jalal et al., 2020) which displays a two phase exponential decay (Fig. S1C). These two unbinding rates may be explained by the small size of the closed DNA probe since, at the beginning of the decay measurement, the DNA probe is saturated with one ParB every 16-bp (see note in Fig. S3) which might lead to an initial fast unbinding and to a slow one when the probe becomes unsaturated. We used the lower rate (~0.017 s^−1^) in the modeling (Fig. 2A) since *in vivo* and in the absence of roadblocks, ParB diffuses rapidly on large DNA molecules, plasmids or chromosomes. However, to better fit the ChIP-seq data, the unbinding rate has to be set four-times lower (Fig. 2B). This optimized value corresponds to ParB clamps opening after ~3.6 min on average. It was previously shown that ParBs are highly dynamic with a residence time inside the clusters of ~100 sec, *i.e.* an exchange rate of 0.01 s^−1^ (Debaugny et al., 2018; Guilhas et al., 2020), a value in favor of the *in vitro* estimate (Fig. S1C and Jalal et al., 2020).

Although the values for these three parameters sound biologically relevant, it is important to note that our modeling is performed on ideal DNA, *i.e.* on a naked DNA without any protein bound to it. The ParB clamp harbors a central hole enabling the DNA molecule to pass through but not protein-DNA complexes (Soh et al., 2019). The unidimensional diffusion on a filament is a physical process that is completely interrupted by roadblocks such as any protein bound stably to the DNA. The ‘Clamping & sliding’ model predicts that ParB clamps would accumulate rapidly between a roadblock and *parS* leading to a plateau, saturated or not, depending on the value, high or low, of the unbinding rate used (Fig. S2). Strikingly, this result is in stark contrast with the nearly symmetrical decreasing pattern observed on both sides of *parS* up to the roadblock (Fig. 2B). On bacterial genomes, with an average density of one gene every Kb and with numerous transcriptional regulators bound to promoter regions (Browning and Busby, 2004), the probability of having obstacles with finite residency times over the > 15-Kb of ParB binding pattern around *parS* is very high. However, we expect these obstacles to modify only weakly and locally the average expected binding profile, so we neglected them in the simulations to focus on the main physical aspects. Therefore, it is very unlikely that numerous ParB could cover a large genomic distance by free diffusion without being halted many times before unloading from DNA. With a ‘Clamping & sliding’ scenario, a higher density of ParB close to *parS* with an important decrease at each locus bound by a protein is rather expected, but never observed in ChIP-seq data (e.g. Baek et al., 2014; Breier and Grossman, 2007; Debaugny et al., 2018; Donczew et al., 2016; Lagage et al., 2016; Sanchez et al., 2015). Also, dips in the ParB DNA binding pattern corresponding to promoter regions with transcriptional regulator binding sites have been previously described both on plasmids and chromosome DNA (Debaugny et al., 2018). Notably, a strong dip in the ParB binding pattern was observed ~1-Kb on the right side of *parS*_F_ inserted on the *E. coli* chromosome, corresponding to the presence of a promoter in reverse orientation relative to ParB diffusion. This suggests that transcription prevents the diffusion of ParB clamps. However, after this important decrease in intensity, the ParB DNA binding signal fully recovered. Such a behavior with dips and peaks is incompatible with a sole sliding mechanism over large genomic distances.

Partition complexes are in a stationary state most of the time involving > 90% of ParB, but not at the onset of *parS* replication. By fast time-lapse microscopy of fluorescently-tagged ParB, we were able to observe some splitting events corresponding to the plasmid segregation step, and to detect variations of fluorescent intensity that might correspond to the replication of *parS*_F_: a rapid drop in the foci intensity followed by its progressive increase to a higher level than the initial one. This temporal pattern might account for the fast disassembly of partition complexes followed by their progressive reassembly, respectively (Fig. 3B). From our measurements, it takes between 20 and 50 sec to reassemble partition complexes (Fig. 3B and S3). The rate obtained from the *in vitro* data with only one ParB release every 10 sec (RCcre ~ 0.1 s-1; Fig. S1B and Jalal et al., 2020) would be much too slow to account for the reassembly with only 2 to 5 ParB loaded in this period of time (requiring 40 min to load 250 ParB).

A fundamental difference between the ‘Clamping & sliding’ model and previous ParB assembly models is the oriented flux of ParB, making the system out-of-equilibrium. Indeed, these previous models explain the formation of ParB*S* assembly in the framework of thermodynamic equilibrium: ‘1D-spreading’, ‘Spreading & bridging’ in the strong coupling limit, ‘Looping & clustering’ and ‘Nucleation & caging’ (Broedersz et al., 2014; Debaugny et al., 2018; Sanchez et al., 2015; Walter et al., 2018). This notable difference comes from the fact that, in the ‘Clamping & sliding’ model, ParB can only be loaded at *parS*, giving rise to a flux of ParB from *parS* to genomic regions away from *parS*. This leads to a severe limitation for this model: ParB would have to cover long genomic distances by unidimensional diffusion to account for the coverage observed in ChIP-seq experiments. On the contrary, models at thermodynamic equilibrium are based on the exchange of ParB with the cytoplasm (playing the role of a reservoir of ParB) at each genomic coordinate based on ParB-DNA and ParB-ParB interactions. Thus, in the case of equilibrium systems, the enrichment around *parS* is due to favorable energetic interactions (thus increasing the probability of the corresponding microstates) and not to the oriented gradient of ParB from to the unique source as in the ‘Clamping & sliding’ model. These ParB-DNA and ParB-ParB stochastic interactions (Fisher et al., 2017; Sanchez et al., 2015) could give rise to droplet formation (via phase transition), which is a mechanism known to quickly create a high concentration region at a targeted cellular location without requiring a membrane (Hyman et al., 2014), and which has been recently proposed to occur for partition complex assembly (Guilhas et al., 2020).

In summary, the ‘Clamping & sliding’ model is unable to describe the overall ParB DNA binding pattern with previously experimentally determined parameters and only inadequately with best fitted parameters. Moreover, it does not to account for several main aspects of this assembly: (i) the rapid turnover of ParB between clusters, (ii) the absence of accumulation of ParB in front of a roadblock, and (iii) the recovery of the ParB binding after strong dips in the profile. Also, the presence of numerous proteins bound along the DNA would prevent ParB clamps from diffusing rapidly to large genomic distances from *parS* centromere sites. For these reasons, the ‘Clamping & sliding’ model alone is not a plausible physical mechanism for fully explaining the partition complex assembly mechanism. We rather propose the possibility that a combination of two mechanisms is at play for the assembly of higher-order nucleoprotein ParB*S* complexes: one occurring at short distance), namely ‘Clamping & sliding’, and one at long distance. We speculate that the distance covered by diffusing clamped-ParB is of the order of the distance between two genes, i.e. ~1-Kb. Indeed, this distance corresponds to average transcription units that would induce barriers arising from both transcription factors and RNA polymerases for diffusing clamped-ParB. We therefore envision that ParB clamps only accumulate in the close vicinity of *parS* sites. These ParBs have undergone a conformational change, occurring possibly (i) with the transition between the *parS*-bound and the sliding clamp states (Soh et al., 2019) and/or (ii) with the stimulation of CTP hydrolysis switching ParB from the closed (CTP-bound) to the open conformation (apo/CDP-bound) (Osorio-Valeriano et al., 2019), that modify the N-terminal domain involved in ParB-ParB dimer interactions (Osorio-Valeriano et al., 2019; Surtees and Funnell, 1999) rendering these ParB prone to interact with other ParBs. The accumulation of these numerous proned-ParB would increase the number of nucleation points that can further recruit most of the intracellular ParBs into a highly concentrated cluster with a phase transition-like mechanism (Guilhas et al., 2020). The ‘Nucleation & caging’ model remains attractive to explain the ParB-ParB interactions occurring at long-distance (> ~1-Kb) as it currently best describes the ParB DNA binding pattern (Debaugny et al., 2018; Sanchez et al., 2015). Such a combination of two modes of actions, ‘Clamping & sliding’ and ‘Nucleation & caging’ is also compatible with the recent study that reveals the droplet-like behavior of the ParB assemblies (Guilhas et al., 2020). Further experimental and modeling work is needed to provide new insights into this crucial higher-order nucleoprotein complex that drives the segregation of the bacterial DNA.

## Methods

All methods can be found in the accompanying Transparent Methods supplemental file.

## Acknowledgements

We thank all members of the Gedy team for fruitful discussions, and P. Rousseau, B. Ton-Hoang M. Campos and F. Cornet for critical reading of the manuscript. This work was supported by Agence Nationale pour la Recherche (ANR-14-CE09-0025-01), CNRS 80Prime (Numacoiled) grant, CNRS “Modélisation pour le Vivant" Grant (CoilChrom), and by the LabEx NUMEV (ANR 2011-LABX-076) within the I-SITE MUSE.

## Author Contributions

Conceptualization, J.-C.W. and J.-Y.B.; Methodology, J.-C.W. and J.-Y.B.; Software, J.-C.W.; Validation, J.R., J.-C.W. and J.-Y.B.; Formal analysis, N.-O.W., J.P., J.D., F.G., J.-Y.B. and J.-C.W.; Investigation, J.R., J.-C.W. and J.-Y.B.; Resources, J.-Y.B.; Writing – original draft, J.-C.W. and J.-Y.B.; Writing – Review & Editing, J.P., A.P., N.-O.W., J.-C.W. and J.-Y.B.; Visualization, J.-C.W. and J.-Y.B.; Supervision, J.-C.W. and J.-Y.B.; Project administration, J.-C.W. and J.-Y.B.; Funding Acquisition, J.-C.W. and J.-Y.B.

