## Supplemental data for "Physical modeling of a sliding clamp mechanism for the spreading of ParB at short genomic distance from bacterial centromere sites"

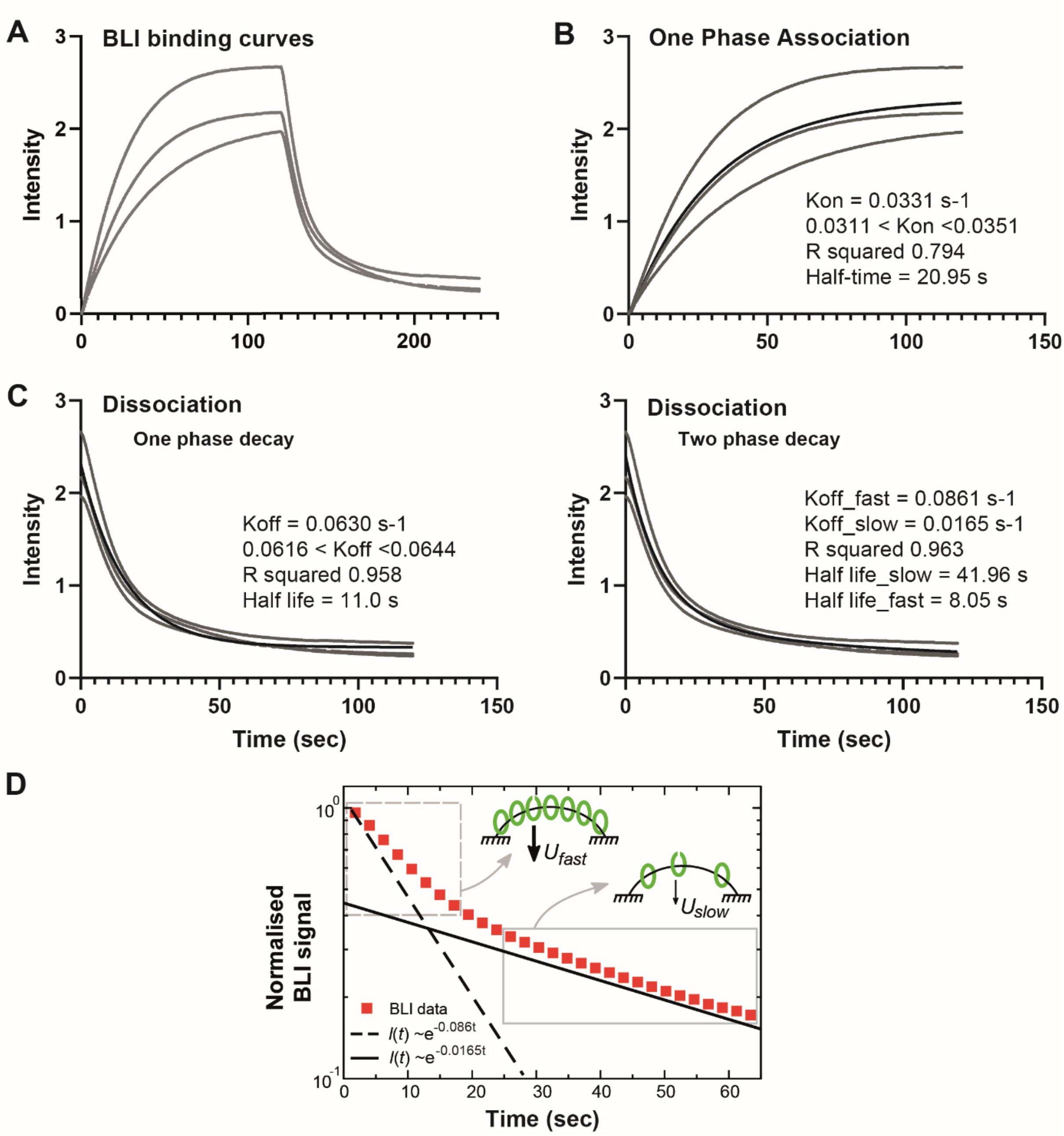


**Figure S1:** Kinetics of ParB loading and unbinding to a closed DNA in the presence of CTP.

The BLI measurement of ParB-DNA interaction was monitored by the wavelength shifts during association or dissociation of ParB to a 169-bp double-stranded *parS* DNA (from Figure 2A in Jalal et al., 2020). The probe DNA is bound at both ends to the sensor surface (no free end). The reactions were measured with and without 1 μM of ParB (dimer) for 120 sec for association and dissociation phases, respectively. The three replicates of the published BLI data were plotted to display the variation in intensity versus time (second). Data points and fitted curves were represented by grey dots and thin black lines, respectively. The parameters computed using nonlinear regression tools (see Methods) were displayed on the right.

**A-** ParB association and dissociation with a closed DNA substrate. The closed DNA containing one *parS* site was infused at time zero with $ParB_{Ccre}$ for 120 sec and then washed out without $ParB_{Ccre}$. Importantly, note that the classical analysis for protein-DNA binding kinetics could not be applied since this model depends on the concentration of ligands. Here, $ParB_{Ccre}$ clamps are delivered from *parS* at a constant rate, i.e. independently of the ParB concentration (ParB concentration is above the KD for ParB-parS binding; Jalal et al., 2020). For this reason, we analyzed independently the association and dissociation phases (see panels **B** and **C**).

**B-** ParB release on non-specific DNA from *parS* analyzed by 'One-phase association' tool. The plot is nicely fitted by an exponential giving a rate of $A \sim 0.033\ s^{-1}$. This rate $A$ corresponds to the ParB release rate from *parS* ($R$) minus the unbinding rate of ParB from DNA ($U$). Therefore, from $A = R - U$, we calculated $R = A + U \sim 0.12\ s^{-1}$ with $U \sim 0.086\ s^{-1}$ (cf panels **C-D**). Note that this experiment, performed in a non-stationary phase starting from an empty DNA, is highly relevant to describe the partition complex recovery stage after DNA duplication upon replication forks passage through *parS*.

**C-** ParB unbinding from DNA. The unbinding curve was analyzed by 'one phase' (left) or 'two phase' (right) 'exponential decay'. The 'one phase' decay provides a Koff $\sim 0.061\ s^{-1}$. The 'two phase decay' provides two dissociation constants: $K_{off\_fast} \sim 0.086\ s^{-1}$ and $K_{off\_slow} \sim 0.017\ s^{-1}$. We used the $K_{off\_fast}$ for estimating the release rate R in panel B since it is performed on the very short and closed DNA probe. However, we used the $K_{off\_slow}$ in our modeling since we propose that it represents the physiological condition where the DNA does not harbor ends attached to a surface close to *parS*. The *in vitro* setup may favor a fast dissociation of ParB from DNA when saturation is reached, a situation that may also occur *in vivo* when ParB accumulates in front of a roadblock on DNA (see main text).

**D-** ParB unbinding from DNA is described by a 'two phase' exponential decay. The signal intensity is normalized to 1 at time 0 and plotted as a function of time in a log-lin scale to discriminate between 'one phase' or 'two phase' decay. Only one replica of the BLI data (red squares) is displayed for clarity. Theoretical curves for exponential decays ($\sim e^{-Koff}$) were plotted with $K_{off} = 0.086\ s^{-1}$ (dotted line) and $0.0165\ s^{-1}$ (black line). This representation clearly indicates that only a 'two phase' exponential decay could fit the ParB unbinding curve. The grey rectangles with dotted and full lines correspond to the data for the fast and slow exponential decays, respectively, and may describe the initial ($U_{fast}$) and steady state ($U_{slow}$) ParB unbinding rates (black arrows), as represented by the corresponding schematic representations (indicated by the grey arrows). Schema represents the in vitro BLI setup with DNA attached at both ends to the surface (black lines). Closed and open green ovals represent ParB dimers in close (clamp) and open confirmations, respectively.

We used the $K_{off\_fast}$ for estimating the release rate R in panel B since it is performed on the very short and closed DNA probe. However, we used the $K_{off\_slow}$ in our modeling since we propose that it represents the physiological condition where the DNA does not harbor ends attached to a surface close to *parS*. The in vitro setup may favor a fast dissociation of ParB from DNA when saturation is reached, a situation that may also occur *in vivo* when ParB accumulates in front of a roadblock on DNA (see main text).

Note: We estimated that the 169-bp DNA could carry ~10 ParB at saturation since ParB dimers could bind every 16-bp without steric hindrance (Sanchez et al., 2015). This estimation is also compatible with the shift of the BLI signal at saturation, which is about 10-times higher than with a unique ParB bound to *parS* (measured in the absence of CTP; Jalal et al., 2020).

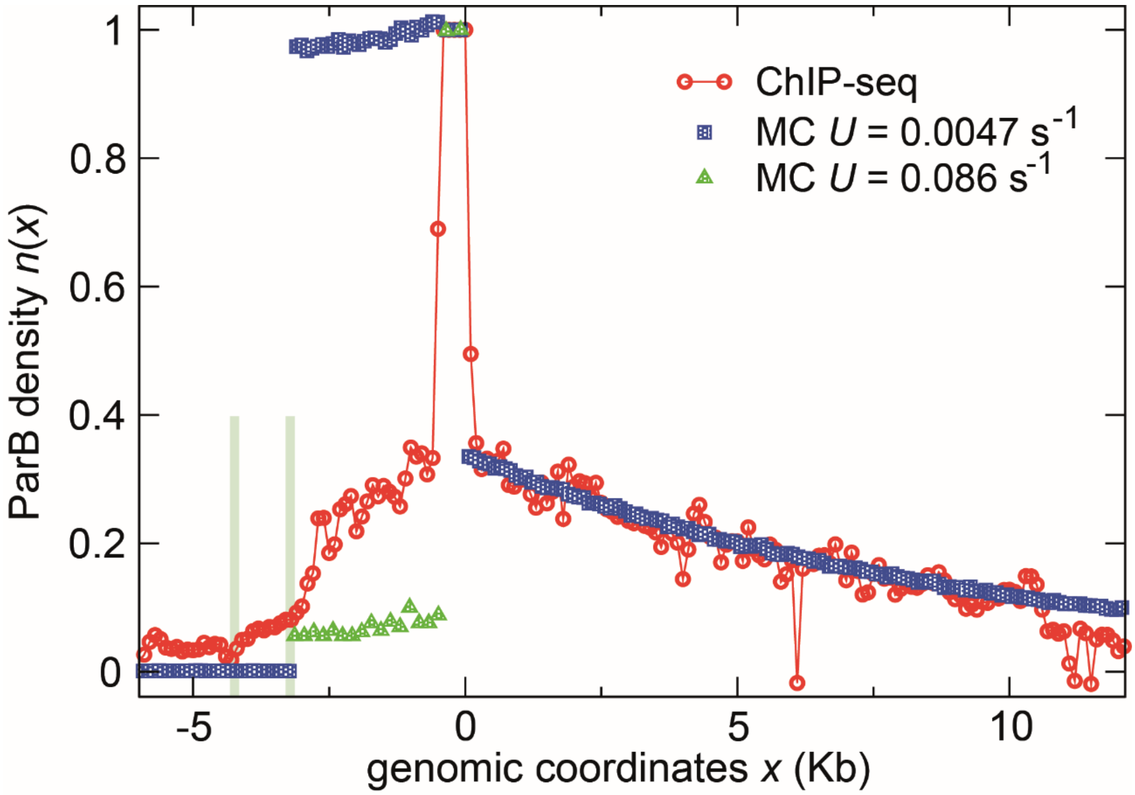


**Figure S2:** The 'Sliding and clamping' model does not describe the ParB binding pattern in the presence of a roadblock.

The ChIP-sequencing data (Sanchez et al., 2015) is represented as in Figure 2B by red circles. The two roadblocks at ~3-kb and 4.5-kb on the left of $parS_F$ are indicated by the green bars. The MC simulations were performed with the indicated values of the unbinding rate $U$. For the higher unbinding rate, the MC simulation (green triangles) is only plotted on the left side of $parS_F$, showing that the decrease also reaches a plateau by contrast to the ChIP-seq data.

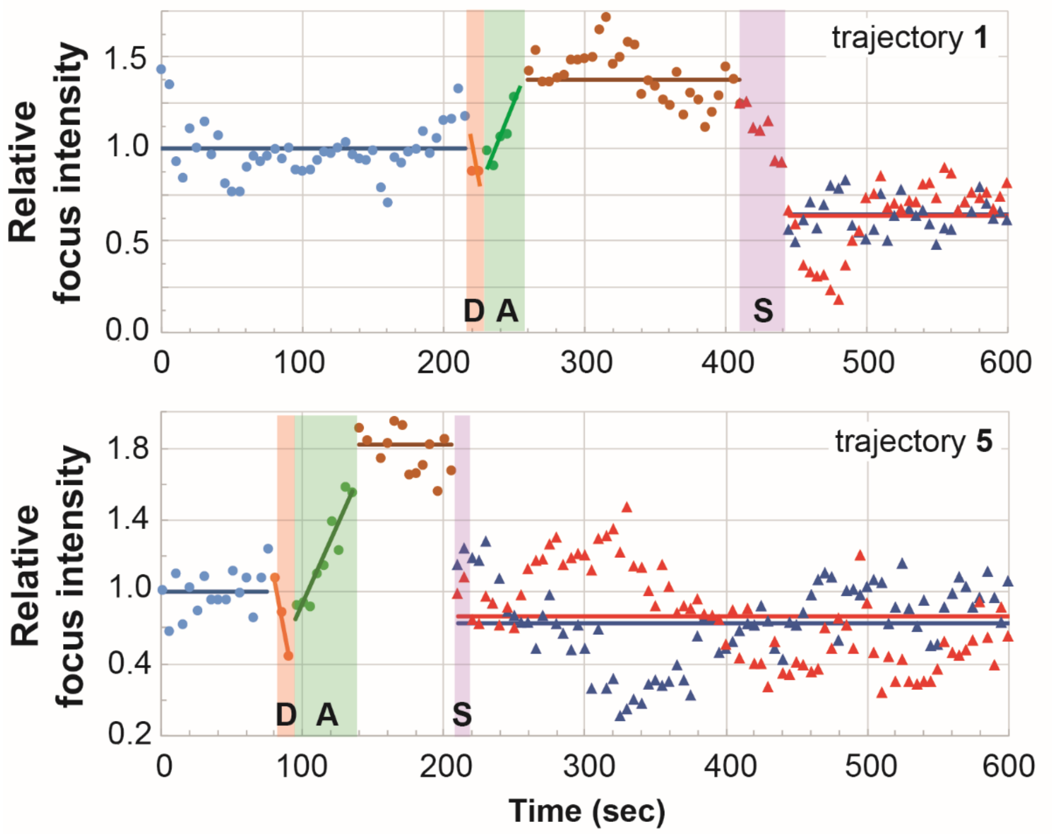


**Figure S3:** Quantification of the fluorescence intensity of the ParB foci over time from Figure 3A. The trajectories number 1 and 5 were quantified and represented as in Figure 3B.

**Table S1:** ParB concentrations inside the cell for *C. crescentus* and the *E. coli* plasmid F.

| | ParB dimers per cell | Mole x $10^{-21}$ | Volume (fL) | Concentration (µM) | *Na* (observed) |
|---|---|---|---|---|---|
| $ParB_F$ | 850 ± 120 | 1.41 | 0.5 | 2.8 | |
| $ParB_F$ (*parS*-associated) | 810 | 1.34 | 0.4 $10^{-4}$ | 10 $10^3$ | ~250 |
| $ParB_F$ (freely diffusing) | 43 | 0.07 | 0.5 | 0.14 | |
| $ParB_{Ccre}$ | 360 ± 40 | 0.6 | 0.25 | 2.4 | |
| $ParB_{Ccre}$ (*parS*-associated) | 290 | 0.48 | 0.2 $10^{-2}$ | 2.4 $10^2$ | ~200 |
| $ParB_{Ccre}$ (freely diffusing) | 70 | 0.12 | 0.25 | 0.48 | |

ParB concentrations are estimated from previous studies that have determined the average intracellular number of $ParB_F$ (Adachi et al., 2006; Bouet et al., 2005 and $ParB_{Ccre}$ (Lim et al., 2014) per cell, the proportion of $ParB_F$ (Sanchez et al., 2015; Guilhas et al., 2020) and $ParB_{Ccre}$ (Lim et al., 2014) in ParB clusters, the number of ParB clusters for the plasmid F (~3.2; Sanchez et al., 2015) and for *C. crescentus* (1 to 2; Lim et al., 2014), and the size (radius) of ParB clusters for the plasmid F (22 nm; Guilhas et al., 2020) and for *C. crescentus* (78 nm; Lim et al., 2014). The volume either corresponds to the volume of the nucleoid over which ParB molecules freely diffuse or to the volume of the ParB clusters for *parS*-associated ParBs. *Na*, the number of ParB clamped on the DNA in the framework of the 'Sliding & clamping model', corresponds to the *parS*-associated ParB divided by the number of cluster per cell.

### Transparent Methods

#### *Monte Carlo simulations*

The simulations data presented in Fig. 2A-B are performed with a sequential Monte Carlo (MC) algorithm. The 60-Kb plasmid is modeled by a filament of $N$ = 3750 sites, each of length $\delta x$ = 16-bp corresponding to the footprint of a ParB protein. The MC time step is defined as a sweep of the $N$ sites of the filament representing the plasmid. When a site is chosen during the MC sweep, different events can occur: if the site is occupied by ParB, this protein (i) can disappear from the lattice at a rate $U$ or (ii) diffuses with a diffusion constant *D;* if the chosen site is neighbor to either side of *parS*, a ParB protein is injected at a rate *R*. We need to perform simulation on a sufficiently small time interval to prevent the appearance of numerical approximations (the analytic solution Eq.(3) are used as a benchmark for the simulations). We used an integration time $\delta t$ corresponding to the time needed for ParB to diffuse over one site (*i.e.* a random step to the right or to the left), thus $\delta t = \delta x^2/D$. The corresponding release rate $R$ and unbinding rate $U$ on the time interval $\delta t$ become $R\delta t = R\delta x^2/D$ and $U\delta t = U\delta x^2/D$, respectively. Thus, we need *D* Monte Carlo iterations to perform an evolution of the system during one second. It is important to remark that, in the absence of interactions between ParB proteins, several particles may be present on the same site. Thus, when a site is chosen, we update all particles on the site.

We define a no-flux boundary condition at *parS*, so that *parS* acts as a barrier for diffusion. Thus a particle is released with a probability one-half on either side of *parS*, *i.e.* the total release rate *R* becomes *R*/2 on each side (this is reflected in Eq.(3)).

In Fig. 2B, the roadblock is defined as an additional barrier located at 3-Kb from $parS_F$ (left side); thus particles that are released on the left side of $parS_F$ have to evolve on an isolated filament of ~3-Kb between the roadblock and *parS*. As the diffusion is fast ($D$ = 4.310$^5$ bp$^2$ s$^{-1}$) with respect to 3-Kb, a good approximation of the distribution is thus a plateau, whose height depends on $\delta x R/(2U \times 3Kb)$. In order to prevent occupancy to become larger than one in the case of large density, we do not allow new particles to be released when the filament is completely covered (*i.e.* saturation).

The initial configuration of the system is empty. Before starting the sampling of the ParB density, we ensure that stationarity is reached. It is helpful to realize that the typical time needed to fill the system at the average stationary value of particles *R/U* corresponds to the release time 1/*R* times *R/U,* thus 1/$U$ corresponds to the average time to fill the system (starting from an empty configuration). It corresponds also to the typical time to replace all the particles of the system in the stationary state, thus it corresponds to the correlation time. In Fig. 2A and 2B, we perform 10 x 1/$U$ seconds of evolution before starting the sampling in order to ensure stationarity. Subsequently, the sampling was started and, to ensure decorrelation of the system between two samplings, samplings were spaced by a time 1/$U$. Averages were performed over 50.000 independent samplings in Fig. 2A and 2B.

We finally note that we can also solve the version of Eq.(2) with a discrete space formulation, i.e. the same framework as Monte Carlo simulations. The discrete solution is the same as Eq.(3) in the limit of $\mu/\delta x \gg 1$, i.e. when the characteristic length of the profile is much larger than the microscopic length $\delta x$ of the system. This limit is satisfied for all the simulations of the paper (~few Kb and $\delta x$ = 16-bp), namely in Fig. 2A where we observed the excellent agreement between both approaches.

The codes used for our simulations are available upon request.

#### *Release (*R*) and unbinding (*U*) parameters*

The kinetic studies for ParB binding and unbinding in the presence of CTP, assayed by bio-layer interferometry (BLI), were performed elsewhere (Fig. 2A in Jalal et al., 2020). To summarize briefly, the measurement of ParB-DNA interaction was monitored by the wavelength shifts during association or dissociation of ParB to a 169-bp biotinylated double-stranded *parS* DNA bound at both ends to the sensor surface (no free end). The reactions were measured with and without 1 µM of ParB (dimer) for 120 sec for association and dissociation phases, respectively.

The uploaded data were analyzed using GraphPad Prism 8© to determine the ParB kinetics parameters. The rates of ParB release from *parS* (*R*) and ParB unbinding from DNA (*U*) were calculated using the nonlinear regression fitting tools 'One phase association' and 'One phase decay' or 'Two phase decay', respectively (Fig. S1).

***ChIP-sequencing data***

High-resolution ChIP-sequencing were from a previous study with $5010^6$ reads per library (Sanchez et al., 2015). Data from the ~60-Kb plasmid F derivative (pOX38B) grown in *E. coli* cells displayed the average number of reads (first nucleotide of each DNA fragment sequenced) per 100-bp windows. The signal is normalized to 1 by averaging the number of reads over the centromere sequence $parS_F$ (550-bp carrying the 1243-bp repeat sequences; Pillet et al., 2011). The drop on the left side corresponds to the RepE/*incC* roadblock (Sanchez et al., 2015).

***Bacterial strain and growth condition***

*E. coli* K-12 strain DLT1215 (Bouet et al., 2006), carrying the reporter mini-F plasmids pJYB249 (Guilhas et al., 2020), were grown at 30°C in M9-Gly (M9 minimal medium supplemented with 0.4% glycerol, 1 mM $MgSO_4$, 0.1 mM $CaCl_2$, 40 µg.ml$^{-1}$ thymine, 20 µg.ml$^{-1}$ leucine and 1 µg.ml$^{-1}$ thiamine) with a generation time of ~100 min allowing the visualization of 1 to 3 plasmids per cell.

***Microscopy and image analyses***

Mid-exponential phase bacterial cultures were sampled, concentrated 5-times by centrifugation and resuspension in M9-Gly, and 0.7 µl was deposited onto slides coated with 1% agarose buffered in M9 solution. Samples were visualized at 30°C as previously described (Diaz et al., 2015), with images taken every 5 seconds over 10 minutes periods. Nis-Elements AR software (Nikon) was used for image capture and editing. Kymographs were generated using the "MultipleKymograph" plugin (ImageJ software). Foci detection and integrated fluorescence were measured using "Trackmate" plugin (Tinevez et al., 2017) in Fiji software.
